# Structural basis of *Toxoplasma gondii* Perforin-Like Protein 1 membrane interaction and activity during egress

**DOI:** 10.1101/399204

**Authors:** Alfredo J. Guerra, Ou Zhang, Constance M. E. Bahr, My-Hang Huynh, James DelProposto, William C. Brown, Zdzislaw Wawrzak, Nicole M. Koropatkin, Vern B. Carruthers

## Abstract

Intracellular pathogens must egress from the host cell to continue their infectious cycle. Apicomplexans are a phylum of intracellular protozoans that have evolved members of the membrane attack complex and perforin (MACPF) family of pore forming proteins to disrupt cellular membranes for traversing cells during tissue migration or egress from a replicative vacuole following intracellular reproduction. Previous work showed that the apicomplexan *Toxoplasma gondii* secretes a perforin-like protein (*Tg*PLP1) that contains a C-terminal Domain (CTD) which is necessary for efficient parasite egress. However, the structural basis for CTD membrane binding and egress competency remained unknown. Here, we present evidence that *Tg*PLP1 CTD prefers binding lipids that are abundant in the inner leaflet of the lipid bilayer. Additionally, solving the high-resolution crystal structure of the *Tg*PLP1 APCβ domain within the CTD reveals an unusual double-layered β-prism fold that resembles only one other protein of known structure. Three direct repeat sequences comprise subdomains, with each constituting a wall of the β-prism fold. One subdomain features a protruding hydrophobic loop with an exposed tryptophan at its tip. Spectrophotometric measurements of intrinsic tryptophan fluorescence are consistent with insertion of the hydrophobic loop into a target membrane. Using CRISPR/Cas9 gene editing we show that parasite strains bearing mutations in the hydrophobic loop, including alanine substitution of the tip tryptophan, are equally deficient in egress as a strain lacking *Tg*PLP1 altogether. Taken together our findings suggest a crucial role for the hydrophobic loop in anchoring *Tg*PLP1 to the membrane to support its cytolytic activity and egress function.

**Author Summary:** *Toxoplasma gondii* has a complex life cycle that involves active invasion of the host cell, the formation of a replicative compartment, and egress from the replicative niche. *T. gondii* encodes a pore-forming protein, *Tg*PLP1, that contains a C-terminal domain that is crucial for efficient exit from both the parasite containing vacuole and the host cell. However, the mechanism by which *Tg*PLP1 recognizes and binds to the appropriate membrane is unclear. Here we use a combination of biochemistry, structural biology, and parasitology to identify the a preference of *Tg*PLP1 for specific lipids and show that a loop within the structure of the C-terminal domain inserts into the membrane and is necessary for egress from the parasite containing vacuole. Our study sheds light into the determinants of membrane binding in *Tg*PLP1 which may inform the overall mechanism of pore formation in similar systems

## Introduction

Cellular egress from the host is a crucial step in the infectious cycle of intracellular pathogens. Accordingly, such pathogens have evolved multiple exit strategies, which can be divided into those that leave the host cell intact and those that rupture the host cell. Several bacterial pathogens, including *L. monocytogenes*, use an actin-based protrusion mechanism that allows a bacterium to enter a neighboring host cell without damaging the original host cell. ^[1]^ Other bacteria have developed extrusion or expulsion mechanisms that also leave the host cell intact. ^[2-5]^ Also, pyroptotic and apoptotic mechanisms leverage cell-death signaling as a means for intracellular pathogens to exit the host cell. ^[6]^ Many apicomplexan parasites, including *Toxoplasma gondii*, use a cytolytic mechanism of egress that obliterates the infected cell. Cytolytic egress results in direct tissue destruction and indirect collateral damage from the ensuing inflammatory response, a hallmark of acute infection by apicomplexan parasites and a key aspect of disease. ^[7]^

The apicomplexan phylum encompasses a variety of parasitic genera important to both human and veterinary health including *Plasmodium*, *Cryptosporidium, Eimeria,* and *Toxoplasma*.^[8-11]^ These parasites contain a unique set of apical secretory organelles, micronemes and rhoptries, which discharge proteins involved in parasite motility, host cell manipulation, and egress.^[12-15]^ *T. gondii* is capable of infecting and replicating asexually within virtually any nucleated cell during the acute stage of infection. Its “lytic cycle” can generally be divided into three steps: invasion where the parasite containing vacuole (parasitophorous vacuole, PV) is formed, intracellular replication, and finally egress from the vacuole. Whereas invasion and intracellular replication have garnered considerable attention, egress remains the least understood component of the lytic cycle.

Efficient egress by *T. gondii* critically relies on micronemal secretion of *Tg*PLP1, a member of the membrane attack complex/perforin (MACPF) protein family ^[15]^. MACPF proteins play central roles in immunity (e.g., perforins and complement proteins), embryonic development (*Drosophila* torso-like and mammalian astrotactins), fungal predation (Oyster mushroom pleurotolysins A/B), and cell traversal or egress by apicomplexan parasites. The domain arrangement for *Tg*PLP1 includes a central MACPF domain that is flanked by an N-terminal domain (NTD) and a C-terminal domain (CTD). Both the NTD and CTD have membrane-binding activity, but only CTD is crucial for proper *Tg*PLP1 function.^[16]^ Although structural insight into the *Tg*PLP1 apicomplexan perforin β-domain (APCβ), which comprises a portion of the CTD, was reported recently,^[17]^ how structural features of this domain contribute to the function of *Tg*PLP1 in egress has not been addressed.

Overall, the mechanism of MACPF proteins begins with membrane recognition by the CTD.^[18]^ Following membrane binding, MACPF proteins oligomerize into ring or arc shaped complexes and undergo a marked structural rearrangement of the MACPF domain where the so-called CH1 and CH2 helices unfurl and become extensions of the central β-sheets to create a large pore.^[19]^ Previous work has shown that *Tg*PLP1 and other apicomplexans share this general mechanism of pore formation. However, the molecular determinants of membrane recognition and binding remain poorly understood. Here, we present evidence that *Tg*PLP1 has a strong preference for inner leaflet lipids and that binding to such membranes occurs via the CTD. Additionally, we solved the 1.13 Å resolution X-ray crystal structure of the TgPLP1 APCβ domain and identified a hydrophobic loop that likely inserts into the target membrane. Finally, we use CRISPR/CAS9 to insert mutations into the hydrophobic loop and show that it is critical for egress competence.

## Results

### PLP1 preferentially binds inner leaflet lipids

Previous work has identified PLP1 as an important egress factor that can bind membranes through its N-terminal domain (NTD) and C-terminal domain (CTD); however, only the CTD has been shown to be required for lytic activity.^[16]^ In the canonical pore forming mechanism, MACPF/CDC proteins bind to the target membrane via the CTD as a first step. Membrane interactions occur via specific protein ^[20-22]^ or lipid ^[23-26]^ receptors in the target membrane. To test if *Tg*PLP1 is capable of binding lipid receptors, we generated liposomes that mimic the outer leaflet (OL) or inner leaflet (IL) of the plasma membrane and tested the binding of both native *Tg*PLP1 and a series of recombinant His-tagged constructs of *Tg*PLP1 **(Figure 1A)** via membrane flotation. Native and recombinant full length *Tg*PLP1 (r*Tg*PLP1) both showed a striking preference for IL liposomes **(Figure 1B & C)**. Recombinant *Tg*PLP1 CTD (rCTD) also showed a preference for IL liposomes whereas the recombinant NTD (rNTD) bound equally to IL and OL liposomes **(Figure 1B & C)**. As a control, we included an unrelated His-tagged recombinant micronemal protein (r*Tg*MIC5), which failed to bind liposomes.

**Figure 1.**
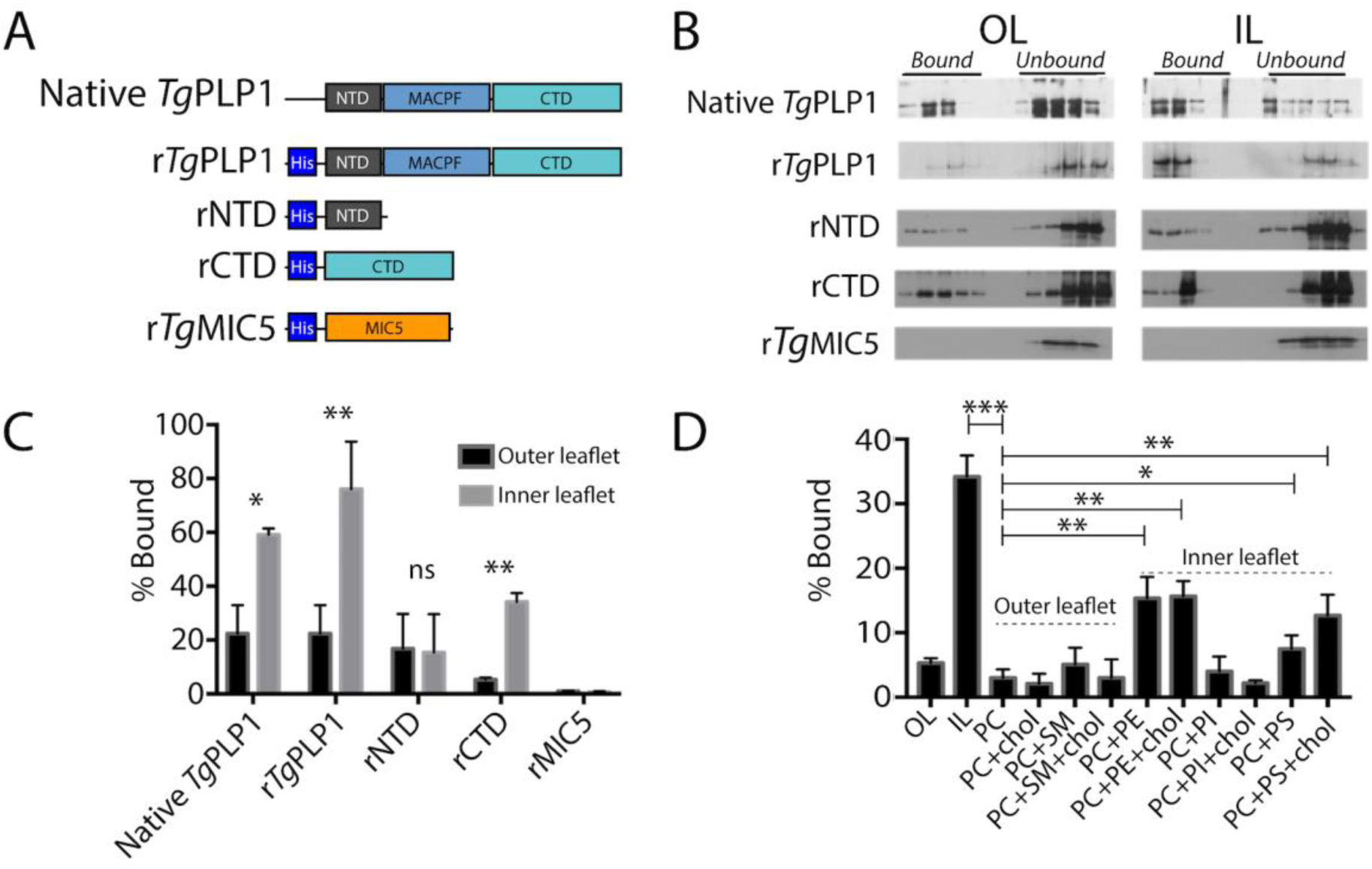
*T. gondii* PLP1 CTD prefers inner leaflet phospholipids. A. Schematic representation of the protein constructs used to test lipid binding. N-terminal, pore-forming, and C-terminal domains (NTD, MACPF, and CTD, respectively) are labeled for clarity. Micronemal protein r*Tg*MIC5 was included as a negative control. B. Representative western blots from membrane flotation assay. After ultracentrifugation aliquots were taken starting at the top of the ultracentrifuge tube. Material contained in the top half of the tube is considered bound to the liposomes while material contained in the bottom portion of the tube is considered unbound. Outer leaflet mimic lipid composition (OL) was as follows: 75% PC, 8.3% SM, 16.7% cholesterol; Inner leaflet mimic lipid composition (IL) was: 35.7% PE, 14.3% PS, 21.4% PI, 28.6% cholesterol. C. Quantification of western blots shown in panel B. Statistical significance determined by paired t-test of three biological replicates. D. Quantification of *Tg*PLP1_CTD_ binding to liposomes of varying composition. Statistical significance determined by unpaired Student’s t-test of three biological replicates.

Since rCTD appears to bind preferentially to liposomes that mimic the IL of the plasma membrane we tested if rCTD prefers binding liposomes composed of individual lipids or combinations thereof. Consistent with our previous observation, rCTD does not bind to liposomes comprised of OL phospholipids, namely phosphatidylcholine (PC) and sphingomyelin (SM), in the presence and absence of cholesterol (**Figure 1D & Figure S1)**. Consistent with its preference for certain IL lipids, rCTD bound to liposomes composed of phosphatidylethanolamine (PE) or phosphatidylserine (PS), but not phosphatidylinositol (PI) **(Figure 1D)**. However, none of the liposomes prepared with individual IL liposomes fully recapitulate the binding to IL mimic liposomes. Together these findings suggest that the preference of *Tg*PLP1 for IL lipids is conferred by its CTD and involves amalgamated binding to PS and PE.

### The TgPLP1 APCβ domain contains a hydrophobic loop that likely inserts into membranes

*Tg*PLP1 contains a well-conserved central MACPF domain that is flanked by a poorly conserved NTD and CTD (**Figure 2A**). The CTD includes the apicomplexan specific APCβ domain, which consists of 3 direct repeats with 4 highly conserved cysteines in each repeat and a C-terminal tail (CTT) that includes a basic patch. To better understand the molecular determinants that govern membrane binding and lipid specificity in this system we expressed, purified, and crystallized the *T. gondii* PLP1 CTD. Despite using a construct that encompassed the entire CTD, the resulting crystals contained only the APCβ domain. Whether this discrepancy is due to a lack of electron density or enzymatic cleavage of the CTT remains unclear.

**Figure 2.**
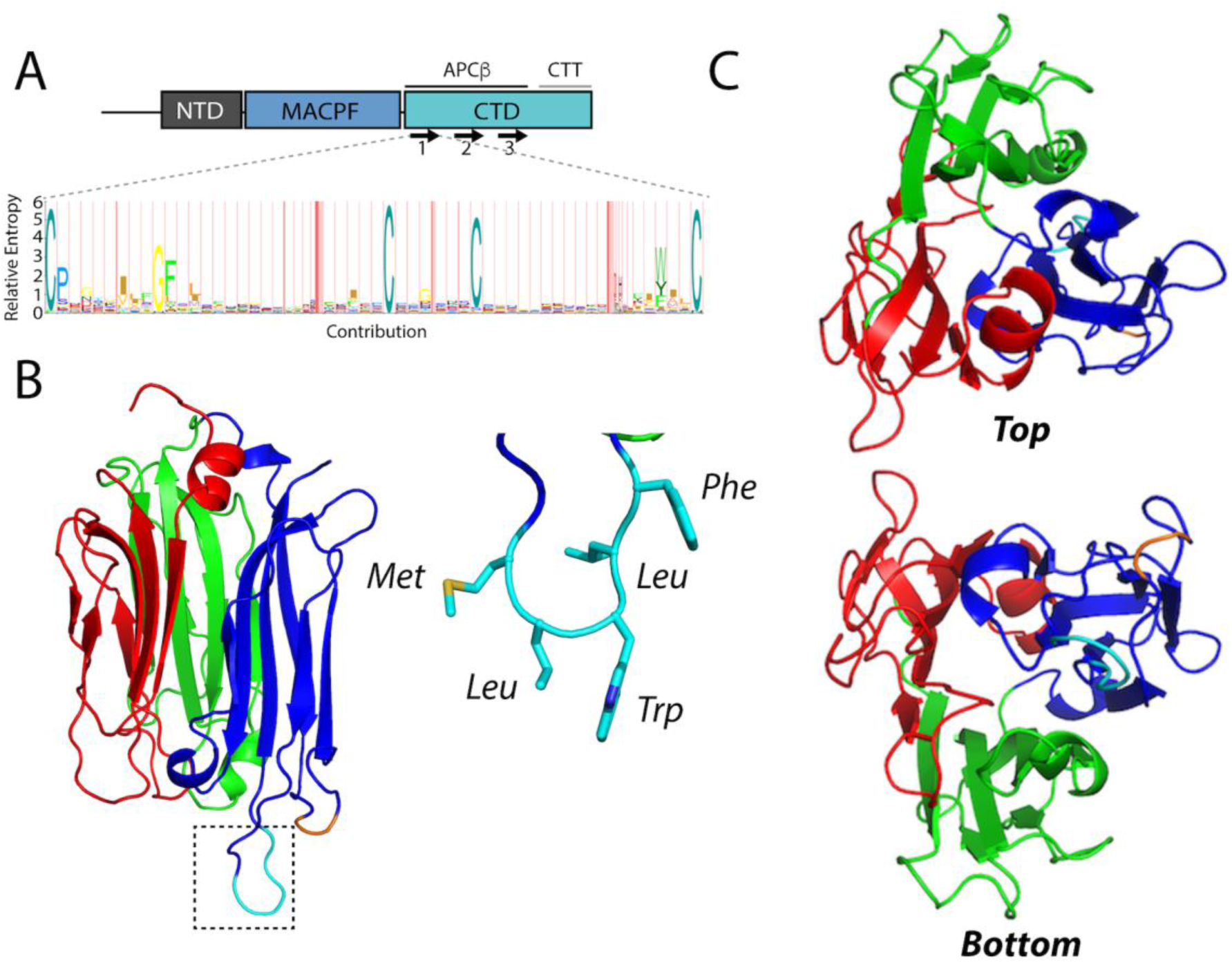
The 1.13 Å resolution crystal structure of the *Tg*PLP1 APCβ domain. A. Schematic representation of the three domains in *Tg*PLP1. The CTD contains a three β-rich sequences (denoted by arrows) and a C-terminal tail (CTT). A Hidden Markov Model consensus display of the β-rich repeat sequences from 51 apicomplexan CTD sequences. B. 1.13 Å resolution crystal structure of the APCβ domain. The crystal structure is made up of three subdomains (red, green, and blue) and contains a basic loop (orange) and a hydrophobic loop (cyan, inset) that protrude from the bottom of one subdomain. C. Top and bottom view of the *Tg*PLP1 APCβ domain highlights the internal pseudo threefold symmetry.

The *Tg*PLP1 APCβ domain crystallized in the C_121_ space group with two monomers in the asymmetric unit. The structure was solved using iodine soaks and single-wavelength anomalous dispersion and subsequently refined to 1.13 Å resolution. The crystallographic data and refinement statistics are summarized in **Table 1**. The *Tg*PLP1 APCβ structure shows that the β-rich repeat region is comprised of a single globular domain with internal pseudo threefold symmetry wherein each β-rich repeat forms a subdomain **(Figure 2B & C).** Each of the three subdomains contains an internal antiparallel β-sheet and an outer β-hairpin (**Figure S2A & B**). The inner and outer layers of each subdomain are held together, in part, by two disulfide bonds between the highly conserved cysteines. An overlay of the three subdomains highlights the similarity in the core of each subdomain. Indeed subdomains 1 and 2 (**Figure S2C** red and green) overlay with an RMSD of 0.728 Å. Subdomain 3 (**Figure S2C** blue), however, aligns less well with subdomain 1 (RMSD 3.207 Å) and subdomain 2 (RMSD 3.175 Å), with the majority of the alignment differences attributed to two loops that protrude from the “bottom” of subdomain 3. The longer of the two loops has hydrophobic character, and thus is termed the hydrophobic loop **(Figure 2B**, colored cyan), whereas the shorter of the two loops has basic character, termed the basic loop (**Figure 2B**, colored orange).

**Table 1.**
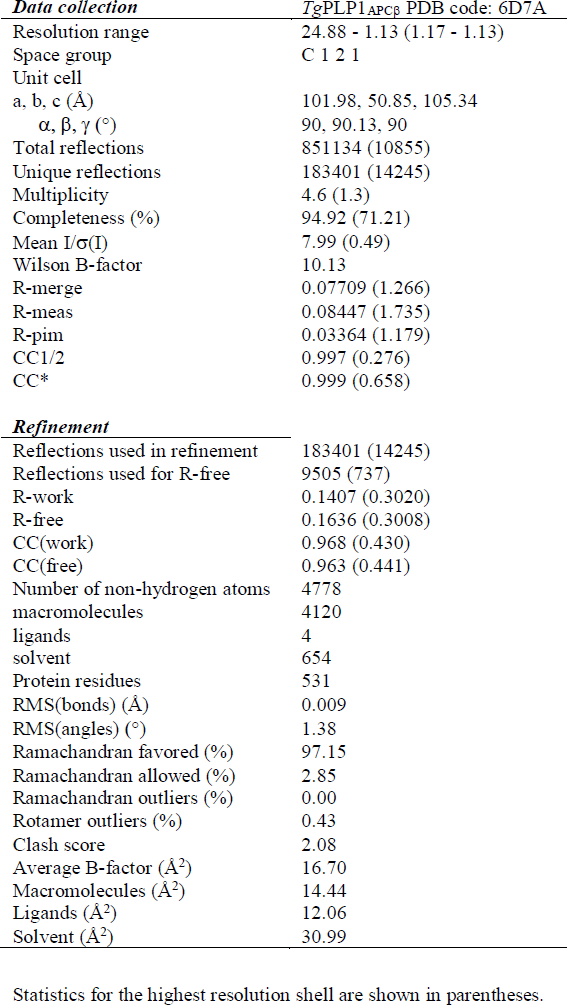
Data collection and refinement statistics

To determine commonality of double-layered β-prism fold we searched for proteins with structural similarity to the APCβ using the Dali server. ^[27]^ The Dali server measures similarity by a sum-of-pairs method that outputs a Dali-Z score. Structures with a Dali-Z score above 2 are considered to have structural similarity. With the exception of a recently published structure of a similar *Tg*PLP1 APCβ construct (Dali-Z score 48.5), ^[17]^ the top scoring result from the Dali server for the *Tg*PLP1_APCβ_ structure is the C-terminal domain of human proprotein convertase subtilisin/kexin type 9 (PCSK9 V-domain, Dali-Z score 6.0; **Table S1**). Indeed, this domain contains similar internal pseudo threefold symmetry and is a double-layered β-prism fold with disulfide bonds that link the inner and outer layers. However, the number of outer layer β-sheets is variable within the domain **(Figure S3)**. Despite the similarities in the overall fold the APCβ structure has very poor structural alignment with the PCSK9 V-domain (RMSD 15.706 Å). These findings suggest that APCβ adopts an uncommon variation of the β-prism fold, which, together with the PCSK9 V-domain, constitutes a new subgroup of the β-prism family.

Given the hydrophobic character of the loop at the bottom of the APCβ structure we reasoned that this loop has the potential to insert into the membrane upon binding. To test this, we took advantage of the intrinsic fluorescence of tryptophan, which is augmented upon exposure to a hydrophobic environment. The *Tg*PLP1 CTD houses four tryptophan residues of which only two are surface exposed **(Figure 3A)**. We recorded the tryptophan fluorescence spectrum of purified rCTD in the presence or absence of liposomes of varying compositions. Incubation with liposomes resulted in an increase in the emission spectrum. Replacement of 10% of PC lipids with PC lipids modified with 1-palmitoyl-2-stearoyl-(7-doxyl)-sn-glycero-3-phosphocholine (7-Doxyl), a collisional quencher, attenuated the increased fluorescence **(Figure 3B & C)**. These observations are consistent with insertion of the hydrophobic loop into the lipid bilayer.

**Figure 3.**
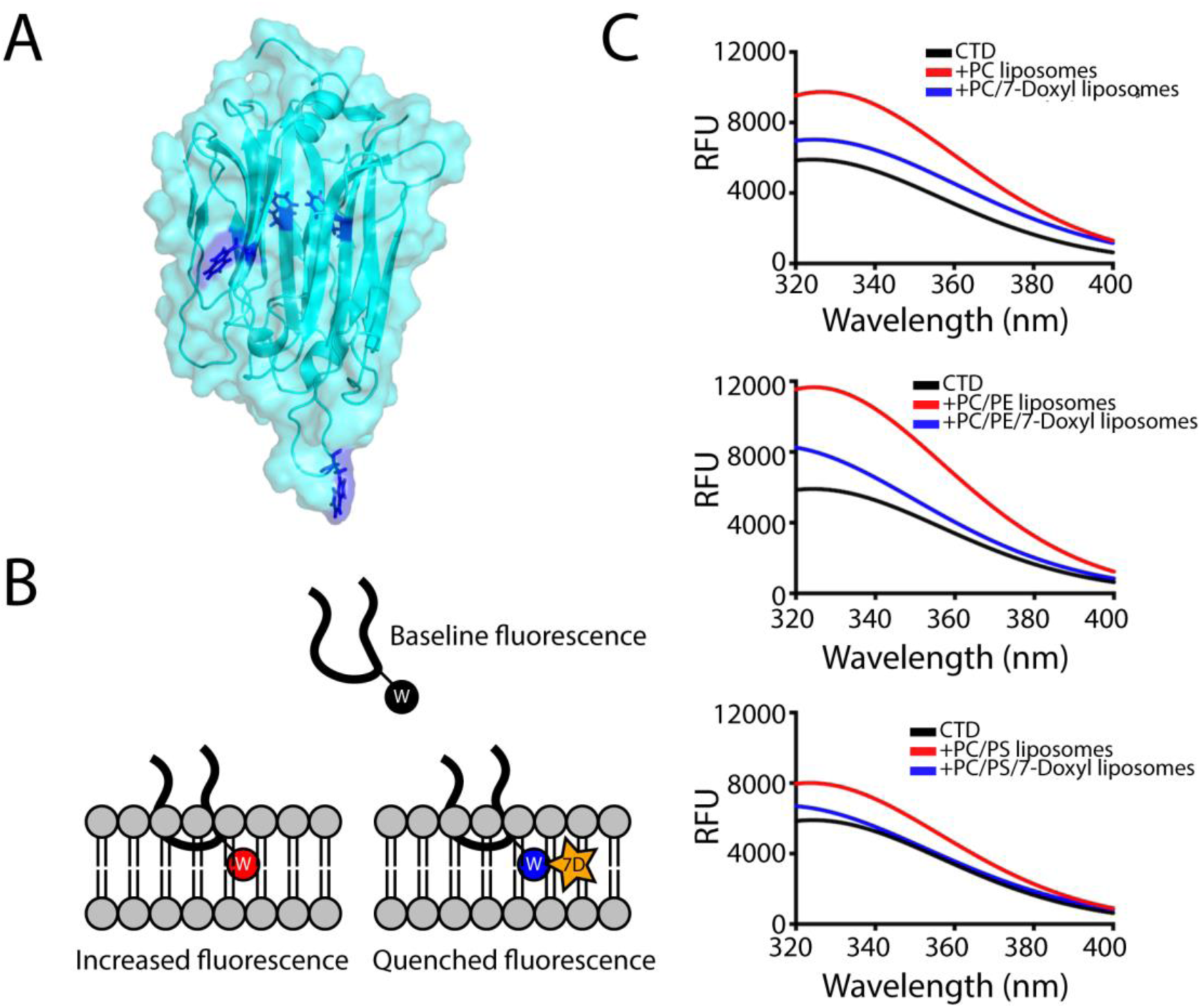
Intrinsic tryptophan fluorescence spectra are indicative of insertion of tryptophan into liposomes. A. Location of the tryptophan side chains in the *Tg*PLP1_APCβ_ structure. Only two tryptophan residues are exposed at the surface of *Tg*PLP1_APCβ_. B. Cartoon schematic of the hydrophobic loop inserting into a lipid bilayer. In the tryptophan fluorescence experiment the intrinsic fluorescence of the tryptophan sidechain increases when moving from an aqueous environment to a more hydrophobic environment in the lipid bilayer. Liposomes were also prepared replacing 10% of the PC lipids with a PC lipid conjugated to a collisional quencher (7-Doxyl, 7D). This results in a quenching of the fluorescence. C. Intrinsic tryptophan fluorescence emission spectra of *Tg*PLP1_CTD_ in the absence (black curves) or presence of PC, PC/PE, or PC/PS liposomes (red curves). In each case the fluorescence emission increases in the presence of liposomes. The increase is less pronounced in the presence of 7-Doxyl (blue curves). Curves are from the averages of three biological replicates each with 3 technical replicates. Liposome composition as follows: PC: 100% PC; PC/7-Doxyl: 90%PC, 10% 7-Doxyl; PC/PE: 50% PC, 50% PE; PC/PE/7-Doxyl: 40% PC, 50% PE, 10% 7-Doxyl; PC/PS: 50% PC, 50% PS; PC/PS/7-Doxyl: 40% PC, 50% PE, 10% 7-Doxyl.

### Integrity of the hydrophobic loop is necessary for parasite egress

We next investigated the importance of the hydrophobic loop in *Tg*PLP1 function. We used CRISPR/CAS9 gene editing to generate mutant parasites expressing *Tg*PLPl with a loop that is symmetrically shortened by two or four amino acids (PLP1_MWF_ and PLP1_W_, respectively) as well as a mutant lacking the loop entirely (PLP1_Δloop_)**(Figure 4A)**. Immunofluorescence microscopy confirmed the micronemal sub-cellular localization of the hydrophobic loop deletion mutants **(Figure S4)**. Additionally, all three loop mutants are secreted in a calcium-dependent manner as evidenced by excreted/secreted antigen (ESA) assays conducted in the presence or absence of the calcium chelator BAPTA-AM **(Figure 4B)**.

**Figure 4.**
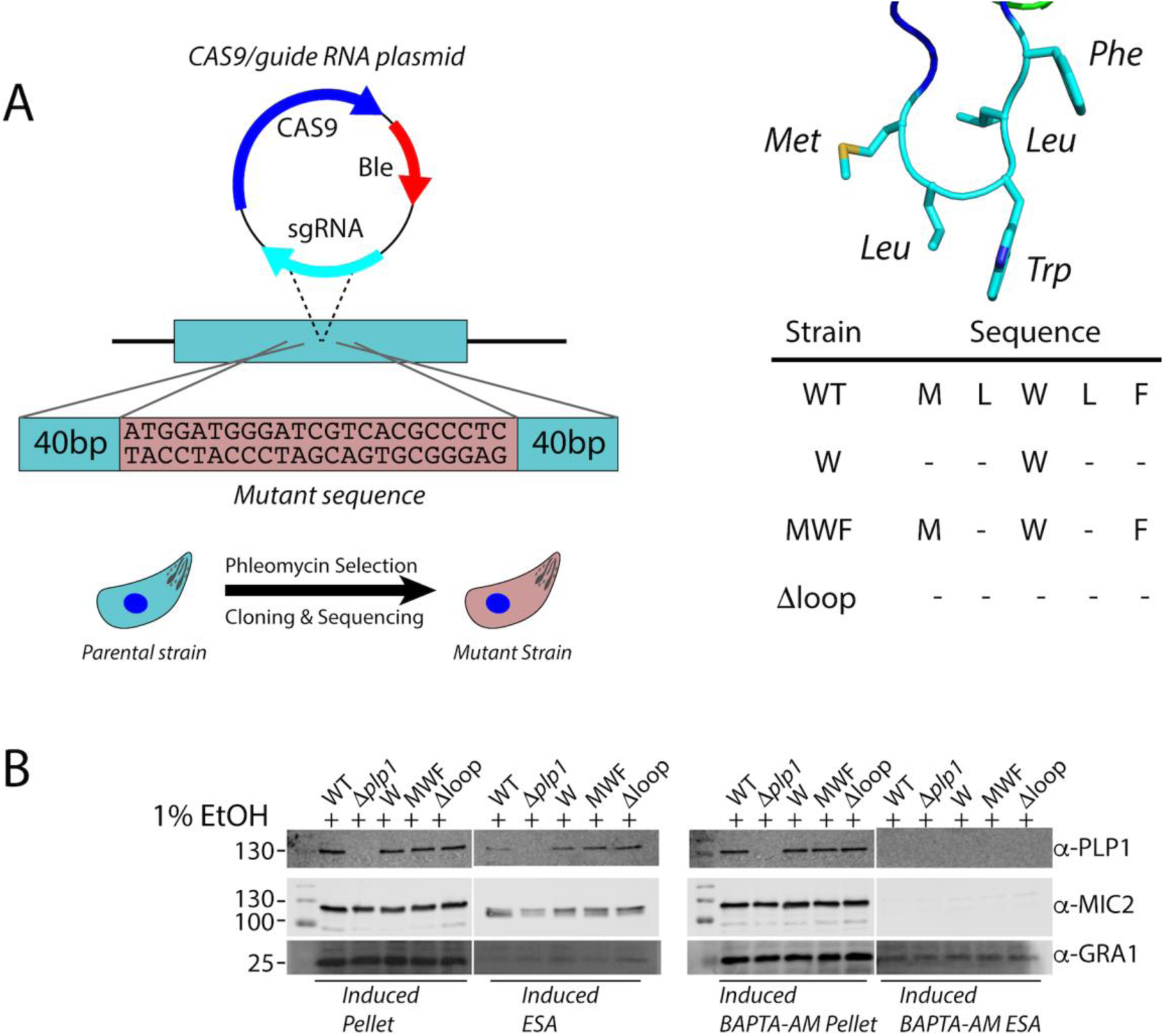
Generation of amino acid deletion mutants in the PLP1 hydrophobic loop. A. Schematic representation of the implementation of CRISPR/CAS9 to generate the mutant strains in this study. A sgRNA was used to generate a double stranded break at the hydrophobic loop and a repair template that encoded the appropriate mutation was supplied. The wild type sequence (MLWLF) was truncated to generate shorter loops with the sequences shown on the right. B. Representative western blot of an ESA assay of amino acid deletions in the hydrophobic loop of *Tg*PLP1, strains are labeled with remaining amino acid sequence as above. Pellet fractions are comprised of parasites recovered after induction. ESA fractions include soluble proteins that have been secreted by the parasites. All strains secrete PLP1 in a calcium dependent manner after induction with 1% ethanol. MIC2 and GRA1 serve as loading controls.

Next, we tested the egress competence of these loop deletion mutants using four general criteria that have been seen in the *Tg*PLP1 knockout strain (Δ*plp1*): (1) the presence of spherical structures in egressed cultures representing failed egress events; (2) formation of smaller plaques relative to wild type parasites; (3) inability to permeabilize the PV membrane (PVM) after calcium ionophore induction; and (4) delayed egress after calcium ionophore induction. Spherical structures were observed in egressed cultures of PLP1_W_, PLP1_MWF_, and PLP1_Δloop_ similar with those seen in Δ*plp1* **(Figure 5A)**. PLP1_W_, PLP1_MWF_, and PLP1_Δloop_ parasites form smaller plaques compared to the parental (WT) strain, consistent to those observed for Δ*plp1* **(Figure 5B & C)**. Previous studies have shown that Δ*plp1* parasites immobilized with cytochalasin D treatment fail to permeabilize the PVM after induced egress with a calcium ionophore as compared to WT parasites. We therefore tested the ability of the hydrophobic loop deletion strains to permeabilize the PVM under the same conditions. PLP1_W_, PLP1_MWF_, and PLP1_Δloop_ parasites all fail to permeabilize the PVM after the addition of 200 μM zaprinast, a phosphodiesterase inhibitor that activates the parasite protein kinase G to induce egress, consistent with what is observed for Δ*plp1* parasites **(Figure 5D)**. Finally, we monitored the extent to which deletions to the hydrophobic loop affect egress from the host cell. To address this, we infected HFF monolayers in a 96-well plate with WT, *Δplp1*, and each of the loop mutants. Thirty hours post-infection the infected monolayers were treated with zaprinast to induce egress. Culture supernatants were assayed for lactate dehydrogenase (LDH), which is released from infected host cells upon parasite egress. *Δplp1*, PLP1_W_, PLP1_MWF_, and PLP1_Δloop_ showed a marked decrease in LDH release compared to cells infected with WT parasites **(Figure 5E)**. These data suggest that integrity of the hydrophobic loop is critical for *Tg*PLP1 function.

**Figure 5.**
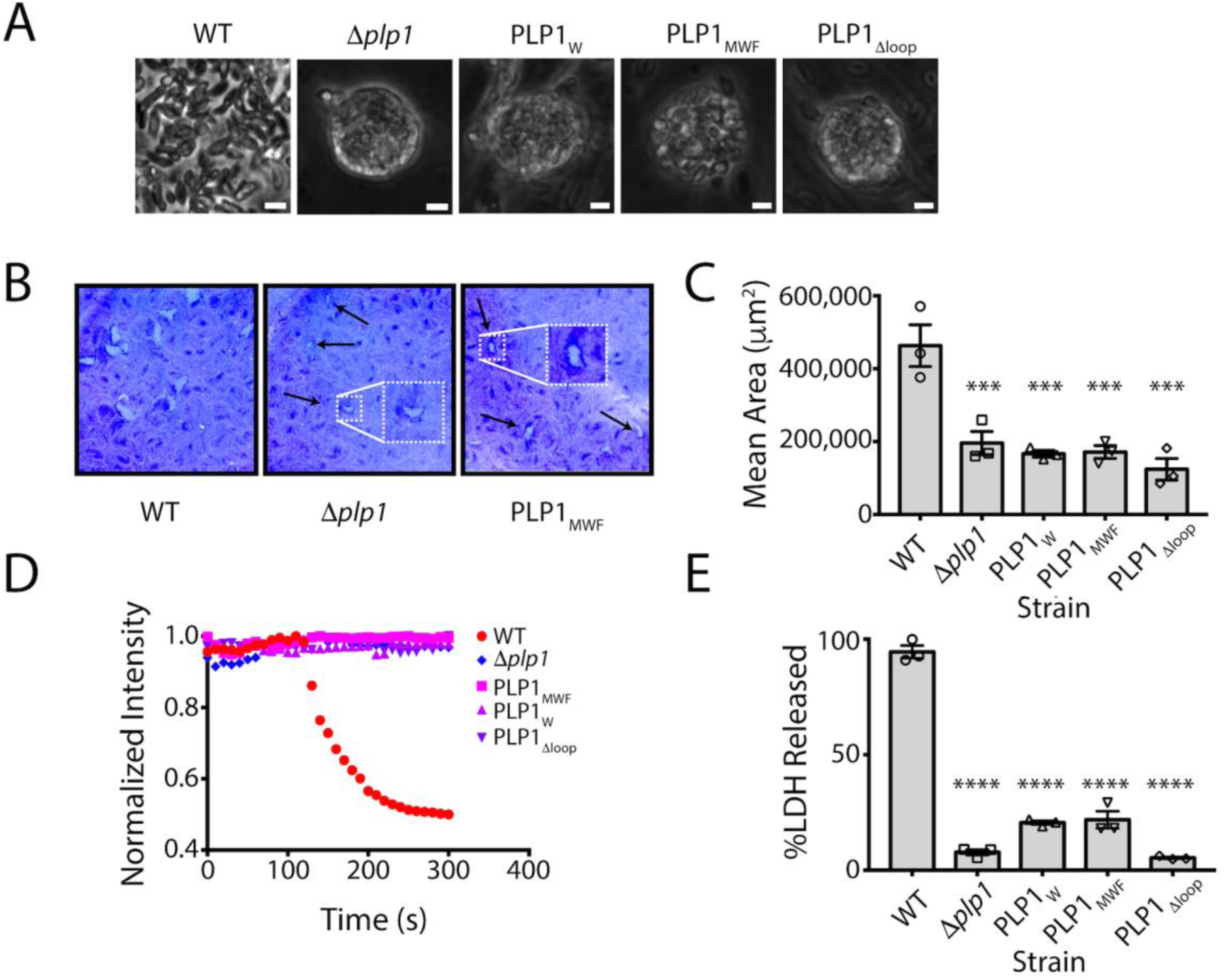
Shortening of the hydrophobic loop mimics the egress phenotype of the *plp1* knockout strain. A. Phase contrast images of parasite cultures. Scale bar, 5 µm. B. Representative images from one of three biological replicate plaque assays for WT, *Δplp1*, and hydrophobic loop amino acid deletion mutants C. Quantification of plaque area. Statistical significance determined by one-way ANOVA with Dunnett correction of three biological replicates. D. Fluorescence intensity tracings of DsRed escape from the PV. Infected monolayers were treated with 1 µM cytochalasin D and 200 µM zaprinast and observed by fluorescence microscopy. E. Egress of wild type and mutant parasites as measured by LDH release upon addition of 200 uM zaprinast and incubation for 20 min. Statistical significance determined by one-way ANOVA with Dunnett correction of three biological replicates.

### Amino acid identity in the hydrophobic loop is important for TgPLP1 function in egress

Since shortening the hydrophobic loop resulted in an egress defect that mimics Δ*plp1* parasites, we tested how the amino acid composition of the loop influences function. We again used CRISPR/CAS9 to generate four mutant strains including a leucine to valine substitution (PLP1_MLWVF_) as well as three alanine substitution mutants (PLP1_MAAAF_, PLP1_MAWAF_, and PLP1_MLALF_) to probe the importance of the leucine and tryptophan residues in the hydrophobic loop. We then tested the subcellular localization of these constructs by immunofluorescence microscopy and confirmed that all mutant strains have micronemal localization of *Tg*PLP1 **(Figure S5)**. Additionally, all four mutants showed calcium-dependent secretion **(Figure 6A)**. Next, we tested the egress competence of these mutant strains using the same criteria described above. Whereas the most conservative mutant (PLP1_MLWVF_) was not retained within spherical structures, all of the alanine substitution mutants (PLP1_MLALF_, PLP1_MAWAF_, and PLP1_MAAAF_) were entrapped in such structures, consistent with defective egress **(Figure 6B)**. Next, we analyzed the plaque size of the mutants. PLP1_MLWVF_ formed large plaques similar to those of WT parasites **(Figure 6C)**. PLP1_MLALF_, PLP1_MAWAF_, and PLP1_MAAAF_ parasites, however, formed small plaques akin to Δ*plp1* **(Figure 6C & D)**. In PVM permeabilization assays, PLP1_MLWVF_ was the only mutant that retained activity after induction with zaprinast. Neither PLP1_MLALF_, PLP1_MAWAF_, nor PLP1_MAAAF_ parasites permeabilize the PVM after induction, thus resembling the defect observed in *Δplp1*, PLP1_W_, PLP1_MWF_, PLP1_Δloop_ strains **(Figure 6E)**. Finally, we tested the extent to which amino acid substitutions to the hydrophobic loop affect egress from the host cell using the LDH assay described above. Consistent with the above findings, PLP1_MLWVF_ is the only point mutant that retained the ability to egress from host cells. PLP1_MLALF_, PLP1_MAWAF_, PLP1_MAAAF_ failed to egress in the duration of the experiment **(Figure 6F)**. Taken together these results indicate that amino acid identity and hydrophobic character of the loop are important for proper *Tg*PLP1 function.

**Figure 6.**
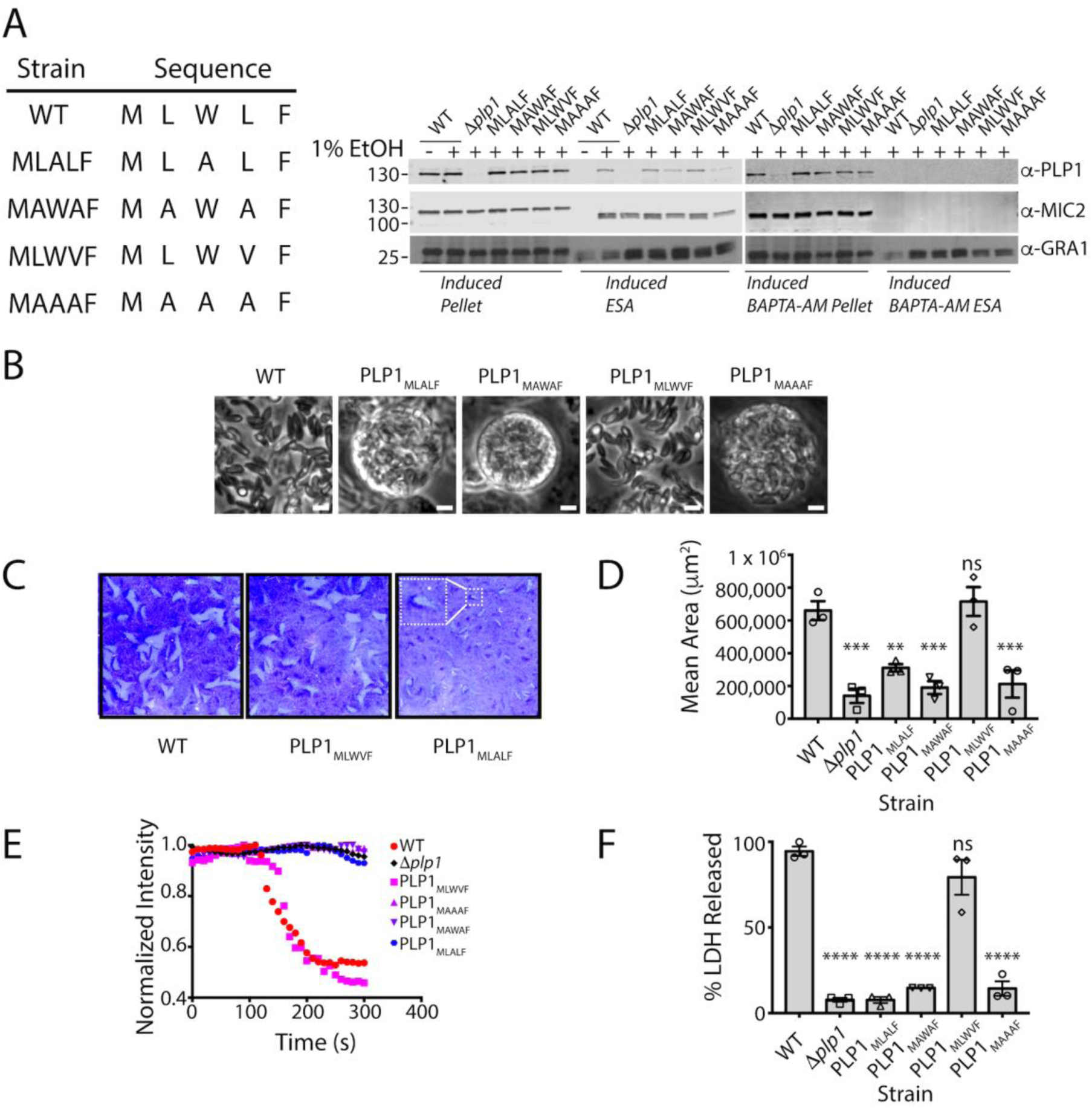
Alanine substitution mutants mimic the deletion strain egress phenotype. A. Representative western blot of an ESA assay of amino acid substitutions in the hydrophobic loop of *Tg*PLP1. The wild type sequence (MLWLF) was mutated to generate various amino acid substitution mutant strains, Strains are labeled with the generated amino acid sequence. Pellet fractions are comprised of parasites recovered after induction. ESA fractions include soluble proteins that have been secreted by the parasites. MIC2 and GRA1 serve as control for calcium-dependent and calcium-independent secretion, respectively. B. Phase contrast images of WT and mutant strains. Scale bar, 5 µm. C. Representative images from triplicate plaque assays of WT, *Δplp1*, and hydrophobic loop amino acid substitution mutants. Inset shows an enlargement of a plaque for a small plaque mutant that exemplifies the other small plaque mutants. D. Quantification of plaque area from three biological replicate experiments. Statistical significance determined by one-way ANOVA with Dunnett correction. E. Fluorescence intensity tracings of DsRed escape from the PV. Infected monolayers were treated with 1 µM cytochalasin D and 200 µM zaprinast and observed by fluorescence microscopy. Data shown is representative of that from three biological replicates F. Egress as measured by LDH release upon addition of 200 uM zaprinast and incubation for 20 min. These data were collected simultaneously with the data shown in Figure 5E. Statistical significance determined by one-way ANOVA with Dunnett correction of three biological replicates.

## Discussion

This paper provides new insight into the structure and function of APCβ, an apicomplexan specific membrane-binding domain associated with parasite egress and cell traversal. Previous studies established that the *Tg*PLP1 CTD, which includes APCβ, has membrane-binding activity and is crucial for *Tg*PLP1 function in egress.^[16]^; however, the lipid binding specificity and structure of this domain remained unknown. The structure solved herein is essentially identical to the *Tg*PLP1 APCβ domain reported recently by Ni et al. ^[17]^, but our work additionally defines the lipid binding specificity, provides evidence for insertion of the hydrophobic loop into membranes, and establishes a critical role for this loop in *Tg*PLP1 function during parasite egress.

We found that the *Tg*PLP1 CTD has a preference for binding IL phospholipids, particularly PS and PE, and that it fails to efficiently recognize OL phospholipids such as PC and SM. These results suggest a working model of directional activity wherein phospholipid accessibility enhances *Tg*PLP1 activity during egress and limits activity during subsequent invasion. In this model, micronemal secretion of *T*gPLP1 during egress delivers it to the PVM for putative binding to PS and PE. Upon escape from the PV, additional secretion of *Tg*PLP1 targets PS and PE in the IL of the host plasma membrane to facilitate parasite exit from the infected cell. The model further posits that a lack of preferred phospholipid receptors on the OL of the target cell limits *Tg*PLP1 activity during invasion, thereby allowing formation of the PV. That *T. gondii* is capable of wounding host cells in a *Tg*PLP1-dependent manner when microneme secretion and *Tg*PLP1 activity are enhanced in low pH medium supports this model. However, the leaflet-specific phospholipid composition of the PVM remains unknown. Although the initial topology of this membrane upon invasion is such that PE and PS would be exclusively in the cytosolic leaflet, reports have suggested extensive lipid remodeling of the PVM after invasion and during intracellular replication. ^[28, 29]^ Also, *T. gondii* is known to secrete into the PV a PS decarboxylase, which converts PS to PE, implying the presence of PS in the PVM ^[30]^. We attempted to create transgenic *T. gondii* that secrete a PS binding probe (lactadherin C2-GFP) into the PV, but the protein is largely retained in the parasite endoplasmic reticulum. An additional limitation of the working model is that the extent to which *Tg*PLP1 acts upon the host plasma membrane during egress remains unknown. Nevertheless, other examples of directional activity exist including the predatory fungus *Pluerotus ostreatus* (Oyster mushroom), which targets nematodes by secreting pleurotolysin A/B, a two-component MACPF pore-forming toxin. The pleurotolysin A subunit requires SM and cholesterol for binding, ^[31-33]^ which is consistent with its action, together with the MACPF B subunit, on exposed membranes of the target nematode. Perforin-1 also exhibits directional activity, but its specificity for a target membrane is influenced by the spacing of phospholipids from unsaturation of acyl chains rather than by recognition of head groups.^[34]^ Defining the phospholipid binding specificity and the role of acyl chain spacing for other MACPF proteins might reveal additional examples of leaflet preference consistent with a directional activity model.

Identifying molecular determinants of membrane recognition by apicomplexan PLPs is necessary to understand cytolytic parasite egress and cell traversal. To address this, we generated an expression construct of the *Tg*PLP1 CTD and solved the 1.13 Å resolution crystal structure of the APCβ domain. Despite using a full-length CTD expression construct the diffraction data only contains the APCβ domain and lacks the CTT. Whether the lack of density is due to flexibility in the CTT or enzymatic cleavage remains unclear. The *Tg*PLP1 _APCβ_ structure presented here is an unusual double-layered β-prism with pseudo threefold symmetry. Analysis of the crystal structure shows a hydrophobic loop that protrudes from one of the subunits, which we interrogated for membrane insertion via fluorescence of a tryptophan located within the loop. The observed increase in intrinsic tryptophan fluorescence in the presence of liposomes is consistent with that seen for perfringolysin O and pneumolysin in the presence of liposomes. ^[35-37]^ The *Tg*PLP1 APCβ domain contains three other tryptophan side chains, but two are buried within the hydrophobic core and their spectra is likely unchanged by the addition of liposomes. The third tryptophan is surface exposed, but is located in between the inner and outer layer of the beta prism, a position that is unlikely to insert into a lipid bilayer. Thus, we conclude that the tryptophan located at the tip of the protruding hydrophobic loop is probably responsible for changes in the fluorescence spectra reported here. These results are consistent with recently published molecular dynamic simulations that theorize insertion of the protruding hydrophobic loop into the membrane. ^[17]^ Insertion of the hydrophobic loop into the target membrane might be required for *Tg*PLP1 to gain a strong foothold for the subsequent conformational rearrangements that accompany pore formation.

To further probe the extent to which the hydrophobic loop is important to *Tg*PLP1 function we generated parasite lines that were genetically engineered to harbor mutations that probed the length of the hydrophobic loop as well as the importance of particular residues therein. Previous work has shown that *Tg*PLP1 is required for efficient lysis of the PVM and the subsequent egress from the host cell.^[15]^ Additionally our previous work has also identified the *Tg*PLP1 CTD as being critical for function.^[16]^ The results presented here show that mutations that affect the length and character of the hydrophobic loop recapitulate the egress defect seen with Δ*plp1* parasites. Namely, these loop mutant strains form spheres in egressed cultures, form small plaques relative to wild type parasites, fail to permeabilize the PVM when motility is arrested, and have a delay in egress after induction. The most striking of these observations is the single alanine substitution of the tryptophan located at the tip of the hydrophobic loop. Tryptophan often contributes to the binding interface of membrane binding proteins because of its properties as both a hydrophobic and polar molecule. Indeed, many MACPF/CDC proteins utilize tryptophan residues for binding target membranes^[35, 38, 39]^ That a single tryptophan to alanine substitution recapitulates the egress defect observed in parasites lacking *Tg*PLP1 altogether underscores the importance of the hydrophobic loop for efficient egress. The extent to which a similarly important loop is conserved in other apicomplexan PLPs remains unknown in the absence of other structural studies.

The work presented here along with that of Ni et al ^[17]^ is a key first step in our understanding of how apicomplexan PLPs recognize and bind to membranes. *T. gondii* encodes two PLPs, *Tg*PLP1 and *Tg*PLP2, but only *Tg*PLP1 has been shown to be important for cytolytic egress. *Tg*PLP2 is not expressed in the tachyzoite stage. A related apicomplexan parasite and causative agent of malaria, *Plasmodium spp.*, has a more complex life cycle and encodes an expanded family of five PLPs, *P*PLP1-5. ^[40-45]^ *Plasmodium* PLPs have been implicated in cytolytic egress as well as in cell traversal, a process required for migrating through tissue. ^[46-48]^ Despite some sequence similarity between the *T. gondii* and *Plasmodium spp.* PLPs (55% similarity between *Tg*PLP1 and *P. falciparum* PLP1) the molecular determinants that govern membrane recognition and binding by *Plasmodium* PLPs remain unclear. More thorough structural and functional studies on *Plasmodium* PLPs are needed to further our understanding of how apicomplexan PLPs recognize their target membranes and shed light on how these differing activities have evolved within a common protein scaffold.

Together with previous findings, the current work brings us closer to a molecular understanding of how *Tg*PLP1 facilitates parasite egress. The work also provides a foundation upon which future work on *Tg*PLP1 and other apicomplexan PLPs are expected to illuminate the molecular basis of differential function in egress and cell traversal.

## Materials and Methods

### Liposome preparation

Liposomes were prepared from 10 mg/mL stock solutions of 1-palmitoyl-2-oleoyl-*sn*-glycero-3-phosphocholine (POPC, Avanti), 1-palmitoyl-2-oleoyl-*sn*-glycero-3-[phospho-L-serine] (POPS, Avanti), L-α-phosphatidylethanolamine (POPE, Avanti), Sphingomyelin (SM, Avanti), L-α-phosphatidylinositol (PI, Avanti), and cholesterol (Avanti) in the ratios described in the figure captions. Two μmol total lipids were dried under a nitrogen stream. Dried lipids were incubated in 1 mL of rehydration buffer (100 mM NaCl, 1 mM CaCl_2_, 1 mM MgCl_2_, 50 mM Tris pH 7.4) for 30 min and subsequently vortexed until lipid film was completely dissolved. Hydrated lipid suspension was subjected to 3 freeze/thaw cycles, alternating between dry ice/ethanol bath and warm water bath. The rehydrated lipid solution was extruded through 400 nm pore-size filters using a mini-extruder (Avanti) to produce liposomes.

### Membrane Flotation

Two hundred nmol of liposomes were incubated with 0.5-1 nmol of recombinant proteins at 37°C for 15 min in a final volume of 200 μL. After incubation the reaction mixture was diluted with 1 mL 85% sucrose and layered with 2.8 mL 0f 65% sucrose and 1 mL of 10% sucrose. The reaction was centrifuged at 115,000g at 4°C for 16 h (Sorvall rotor AH-650). Fractions were collected and analyzed by western blot. Band intensity was quantified using Image J. The liposome binding efficiency was calculated as:

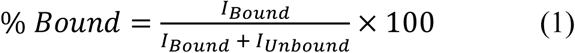

### Protein Production & Purification

High-five or *Sf9* cells were expanded to six 1 L volumes. These were used to seed 20 L of media (Expression Systems) at 2.0 x 10^6^ cells/mL in a 36 L stir tank bioreactor. The culture was infected with recombinant baculovirus at a multiplicity of infection of 5. The reactor was incubated at 27°C with stirring and sparged with air for 72 h. The culture was pumped out and centrifuged at 1000g and 4°C for 40 min to pellet the cells. The media was collected, and batch bound with 0.5 mL per liter of Roche cOmplete His-Tag purification resin for 4 h at 4°C with stirring. Resin was loaded onto a column and washed with 50 mM Tris (pH 8.0), 300 mM NaCl and 20 mM imidazole. Protein was eluted with 50 mM Tris (pH 8.0), 300 mM NaCl and 250 mM imidazole. Wash and elution fractions were collected and run on SDS page to determine purity and protein location. The poly-histidine tag was subsequently cleaved with TEV-protease at 4°C overnight and separated from the CTD by immobilized metal affinity chromatography. CTD fractions were further purified by anion exchange chromatography and concentrated to 22 mg/mL.

### Crystallization & Data Acquisition

*Tg*PLP1_APCβ_ crystals were obtained from the Molecular Dimensions Clear Strategy II screen set up with 1 μL of protein and 1 μL of reservoir containing 100 mM sodium cacodylate at pH 6.5, 0.15 M KSCN, and 18% PEG 3350. Hanging drop plates were equilibrated at 20°C. The needle-like crystals were mounted on a cryoloop and transferred to a cryoprotectant solution containing the reservoir solution supplemented with 20% ethylene glycol. Native crystals were flash frozen in liquid nitrogen for data collection. For phase problem solution data sets, a two-hour soak was performed in a synthetic mother liquor supplemented with 20 mM KI and flash frozen in mother liquor supplemented with 20% ethylene glycol. Data sets were collected at LS-CAT beamline 21ID-D and 21ID-G.

### Structure determination

The crystals belonged to the C_121_ space group with cell dimensions of a = 101.98 Å, b = 50.85 Å, c = 105.34 Å, α = 90°, β = 90.13°, and γ = 90°. Data reduction and scaling were performed with autoPROC.^[49]^ Phasing was performed using the AutoSol in Phenix.^[50]^ Initial solution was obtained by SAD on KI soaked crystals. The partial solution was then used as a molecular replacement model for the high-resolution data sets. Model building was performed in COOT ^[51]^ and refinement in PHENIX. The *Tg*PLP1_APCβ_ crystal structure was refined to a crystallographic R_work_ of 14.07% and a R_free_ of 16.36% The final structure was analyzed with validation tools in MOLPROBITY. Structural visualization was performed via PyMOL.

### Fluorescence Measurements

Intrinsic tryptophan emission intensity was measured on a Biotek Snergy H1 equipped with monochromators for both excitation and emission. The emission spectra were recorded between 320-400 nm with a step size of 5 nm. The excitation wavelength was set to 280 nm. The emission intensity was recorded in the absence and presence of PC, PC/PE, PC/PS liposomes and liposomes that had 10 mol% of the PC lipid replaced with 1-palmitoyl-2-stearoyl-7-doxyl)-sn-glycero-3-phosphocholine (7-Doxyl) (Avanti).

### Host cell and parasite culture

All cells and parasites were maintained in a humidified incubator at 37°C and 5% CO_2_. Human foreskin fibroblast cells (HFF, ATCC CRL-1634) were maintained in Dulbecco’s modified Eagle’s medium (DMEM) supplemented with 10% Cosmic Calf serum, 20 mM HEPES pH 7.4, 2 mM L-glutamine and 50 μg/mL penicillin/streptomycin. All *T. gondii* parasites were maintained by serial passaging in HFF cells and were routinely checked for mycoplasma contamination.

### Parasite line generation

PLP1 hydrophobic mutant strains were generated using CRISPR-Cas9 using RH*Δku80* as a parental strain. 20 bp of *plp1*-specific guide RNA sequence targeting the hydrophobic loop was inserted into pCRISPR-Cas-9-Ble using site-directed mutagenesis to generate the pAG1 plasmid. An annealed synthetic oligonucleotide repair template pair that encoded the appropriate mutations and a silent mutation to replace the NGG cut site (IDTDNA) was mixed with pAG1 and precipitated by addition of ethanol. Fifty million tachyzoites were transfected by electroporation in a 4 mm gap cuvette using a Bio-Rad Gene Pulser II with an exponential decay program set to 1500 V, 25 μF capacitance and no resistance and immediately added to a confluent HFF monolayer in a T25 flask. Parasites were selected with 50 μg/mL phleomycin 24 h post transfection for 6 h and subsequently added to a confluent HFF monolayer in T25 flask. Parasites were incubated for 36-48 h. Clonal populations were isolated and a 1 kb fragment generated from extracted DNA was sequenced for confirmation of the correct mutation.

### Excreted secreted antigen

Induced excreted secreted antigens (ESA) were performed as previously described. ^[52]^ Briefly, 2 x 10^7^ parasites were incubated at 37°C for 2 min in 1.5 mL Eppendorf tubes with DMEM containing 10 mM HEPES, pH 7.4 and supplemented with 1% ethanol. Parasites were separated by centrifugation (1000g, 10 min, 4°C). Samples of the pellet and supernatant were run on a 10% SDS-PAGE gel and analyzed by western blot. ESAs were performed in the presence and absence of 100 μM BAPTA-AM.

### Immunofluorescence staining

Infected monolayers were fixed with 4% formaldehyde for 20 min and washed with PBS. Slides were with 0.1% Triton X-100 for 10 min and blocked with 10% fetal bovine serum (FBS), 0.01% Triton X-100 in PBS). Slides were subsequently incubated with rabbit anti-PLP1 (1:500) and mouse anit-MIC2 (6D10, 1:250) diluted in wash buffer (1% FBS, 1% Normal Goat Serum (NGS), 0.01%Triton X-100 in PBS) for 1 h at RT. After three washes, slides were incubated for 1 h at RT with Alexa Fluor goat anti-rabbit and goat anti-mouse secondary antibodies (Invitrogen Molecular Probes) diluted in wash buffer (1:1000). Slides were washed and mounted in Mowiol prior to imaging.

### Plaque assay

Infected monolayers of HFF cells in T25 flasks were washed with phosphate-buffered saline (PBS). Parasites were liberated by scraping the monolayer and passaging through a 27-gauge needle. Liberated parasites were filtered through a 3 μm filter (Millipore) and counted on a hemocytometer. Monolayers of HFFs in individual 6-well plates were inoculated with 50 parasites and incubated, undisturbed, for one week and fixed with 2% (w/v) crystal violet. Plaque size was quantified using ImageJ.

### PV permeabilization

Monolayers were infected with tachyzoites expressing DsRed and allowed to replicate for 30 h. Infected monolayers were washed twice with warmed Ringer’s buffer (155 mM NaCl, 3 mM KCl, 2 mM CaCl_2_, 1 mM MgCl_2_, 3 mM,NaH_2_PO_4_, 10 mM HEPES, 10 mM Glucose, 1% FBS, pH 7.40). Ionophore treatment was initiated by addition of an equal volume of 2 μM cytochalasin D and 400 μM zaprinast. Fluorescence image time series were collected every 10 sec for 5 min.

### LDH egress assay

Infected monolayers of HFF cells in T25 flasks were washed with phosphate-buffered saline (PBS) and liberated by scraping the monolayer and passaging through a 27-gauge needle. Liberated parasites were filtered through a 3 μm filter (Millipore), counted on a hemocytometer, and centrifuged at 1000g for 10 min. Parasites were resuspended to a density of 5 x 10^5^ parasites/mL and each well of a 96-well flat bottom culture plate was inoculated with 5 x 10^4^ tachyzoites of the appropriate strain. Infected cells were washed with warm Ringer’s buffer and treated with zaprinast diluted to 200 μM in Ringer’s buffer, Ringer’s with an equal volume of dimethyl sulfoxide (DMSO), or cell lysis reagent (BioVision) diluted in Ringer’s. Treated plates were incubated at 5% CO_2_ and 37°C for 20 min. Plates were placed on ice and 50 μl of each well was transferred to a 96-well round-bottom plate. Round-bottom plates were centrifuged at 500g for 5 min. Ten μl of solution was tested for lactate dehydrogenase (LDH) following the manufacturer’s protocol (Biovision).

## Data availability

The atomic coordinates are available at the Protein Data Bank (PDB) under accession number 6D7A.

## Author Contributions and Notes

Lipid binding experiments were designed by OZ and VBC and performed by OZ. JDP and WCB expressed recombinant proteins and performed initial purification. CMEB purified recombinant proteins and crystallized *Tg*PLP1_APCβ_. Diffraction data was collected by CMEB, NMK, and ZW. The *Tg*PLP1_APCβ_ structure was solved by NMK and refined by AJG. Tryptophan fluorescence and mutational analysis was designed by AJG and VBC and performed by AJG. Mutant characterization was performed by AJG and M-HH. AJG and VBC wrote the manuscript.

The authors declare no conflict of interest.

This article contains supporting information online.

## Acknowledgments

We thank Tracey Schultz for laboratory support and Aric Schultz for expertise and thoughtful discussions regarding *Toxoplasma* egress. We also thank Jingga Inlora, Vineela Chukkapalli, Dishari Mukhejee, Chris Sumner and Akira Ono for help with liposome preparation. We gratefully acknowledge the funding support from the University of Michigan Departments of Chemistry and Biophysics and the U.S. National Institutes of Health (R01AI46675 to VBC). This research used resources of the Advanced Photon Source, a U.S. Department of Energy (DOE) Office of Science User Facility operated for the DOE Office of Science by Argonne National Laboratory under contract no. DE-AC02-06CH11357. Use of the LS-CAT Sector 21 was supported by the Michigan Economic Development Corporation and the Michigan Technology Tri-Corridor (grant 085P1000817).

## Supplementary Figures & Tables

**Figure S1.**
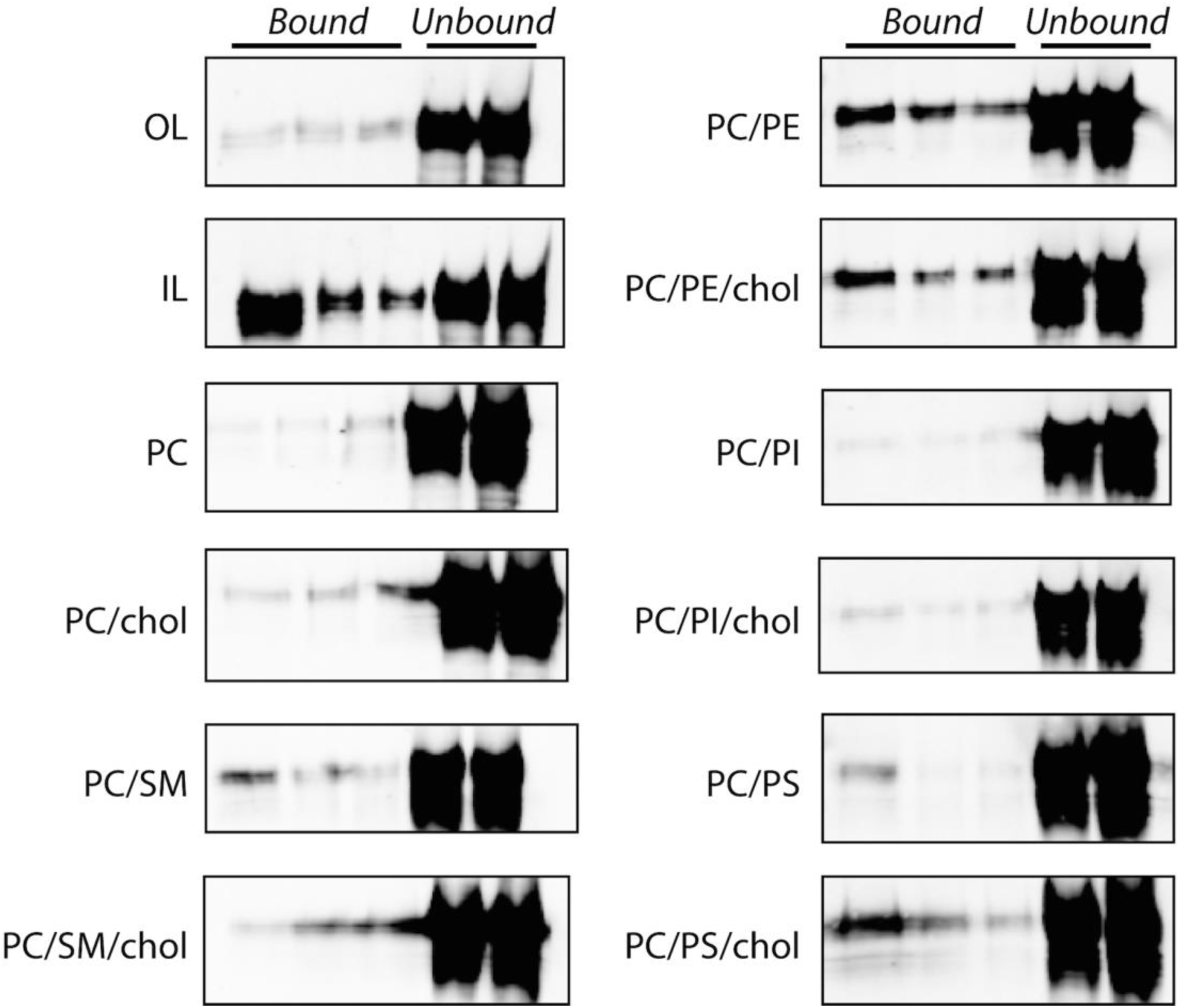
*Tg*PLP1_CTD_ binding to liposomes of varying composition. Representative western blots of liposome membrane flotation assays. Liposome composition as follows: OL: 75% PC, 8.3% SM, 16.7% cholesterol; IL: 35.7% PE, 14.3% PS, 21.4% PI, 28.6% cholesterol; PC: 100% PC; PC/chol: 50% PC, 50% cholesterol; PC/SM: 50% PC, 50% SM; PC/SM/chol: 30% PC, 50% SM, 20% cholesterol; PC/PE: 50% PC, 50% PE; PC/PE/chol: 30% PC, 50% PE, 20% cholesterol; PC/PS:: 50% PC, 50% PS; PC/PS/chol: 30% PC, 50% PS, 20% cholesterol; PC/PI:: 50% PC, 50% PI; PC/PI/chol: 30% PC, 50% PI, 20% cholesterol. Quantification of these bands is shown in Figure 1D.

**Figure S2.**
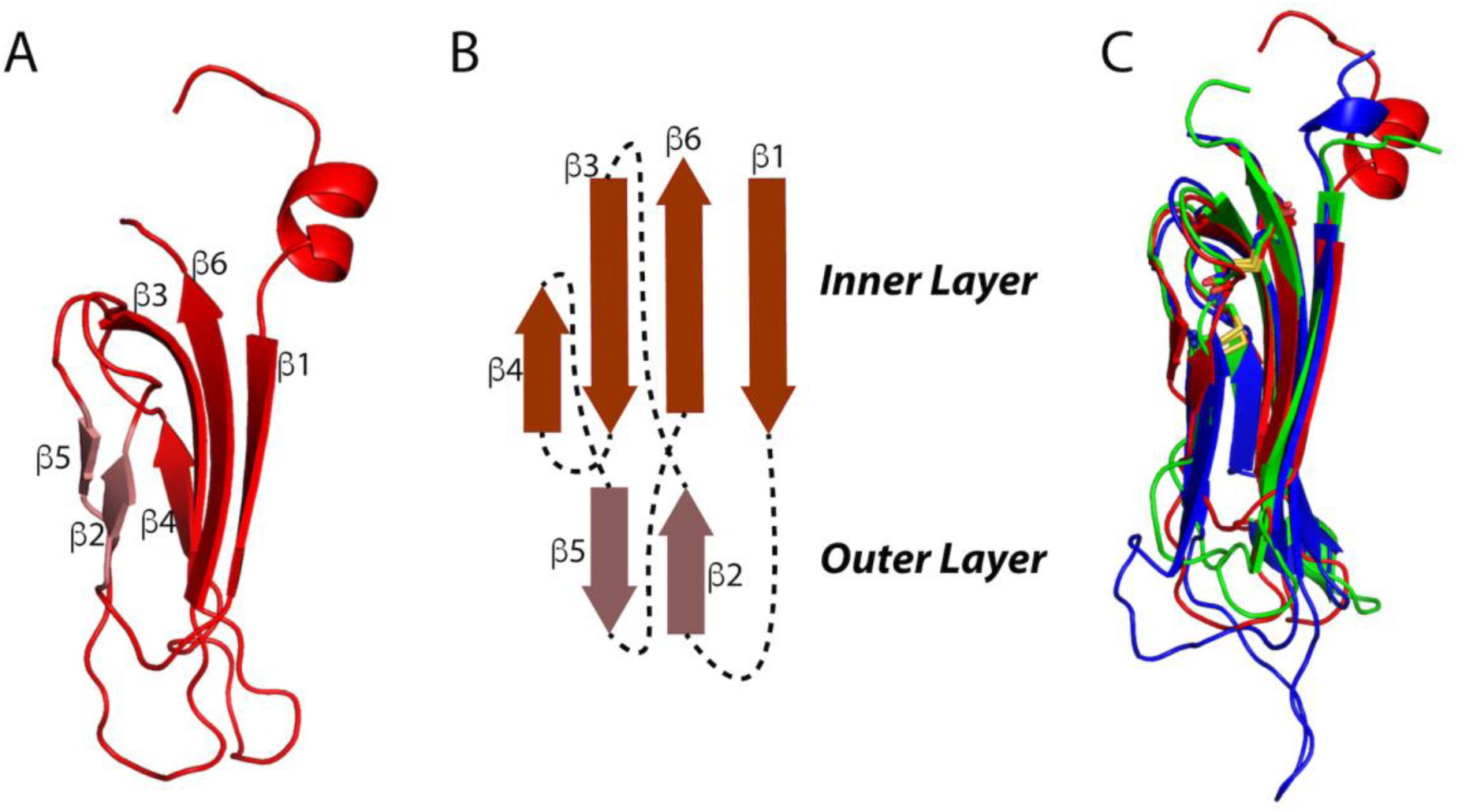
Structural details of the *Tg*PLP1 APCβ structure. A. Subdomain 1 of the *Tg*PLP1_APCβ_ crystal structure. Each subdomain is made up of an inner β-sheet of four anti-parallel strands and an outer β-sheet of two anti-parallel strands. B. Schematic representation of the inner and outer β-sheets. C. Overlay of the three subdomains of the APCβ crystal structure.

**Figure S3.**
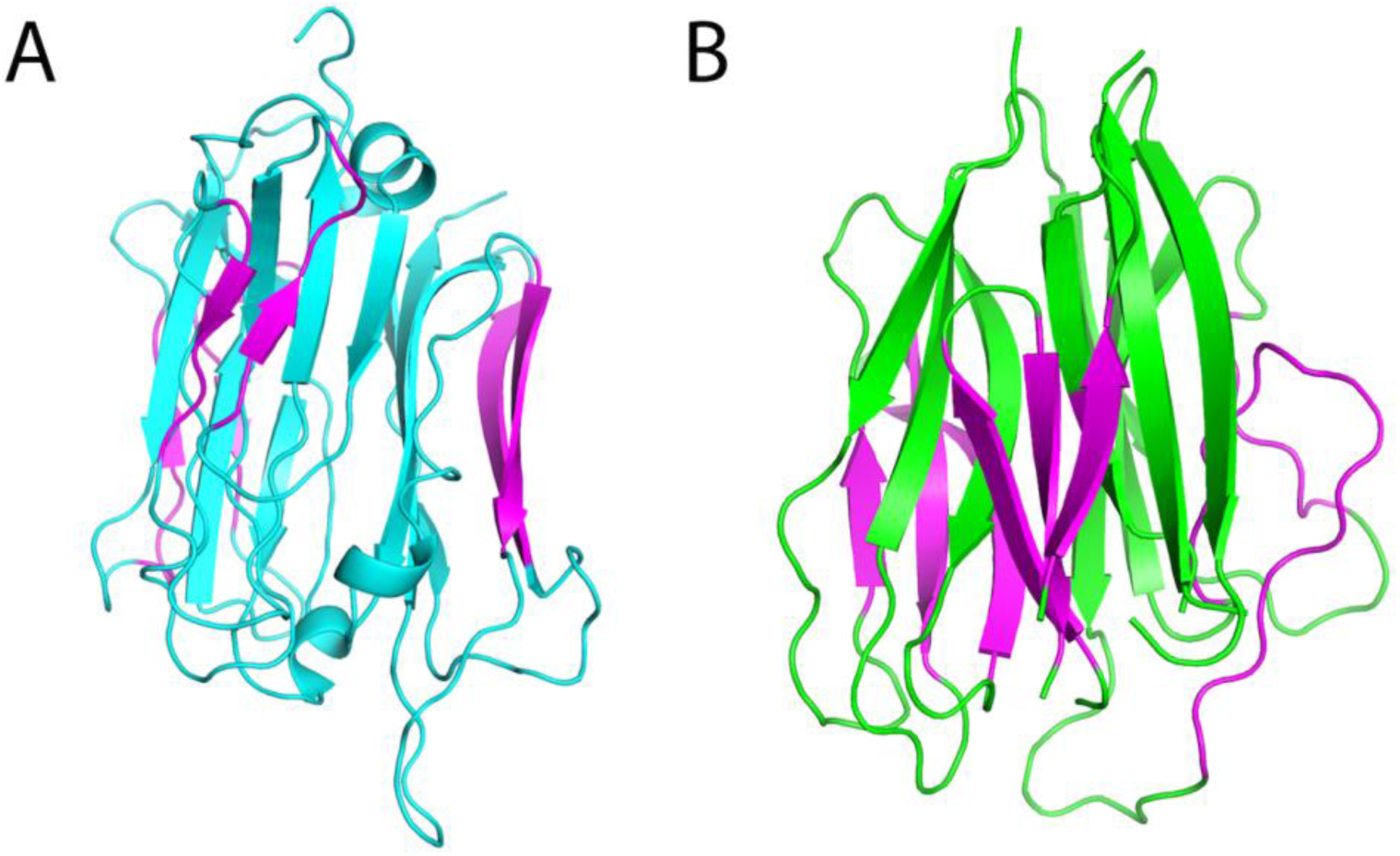
Comparison of the *Tg*PLP1 APCβ crystal structure and the PCSK9 V-domain crystal structure. A. Cartoon representation of *Tg*PLP1 APCβ structure. B. Cartoon representation of the PCSK9 V-domain.

**Figure S4.**
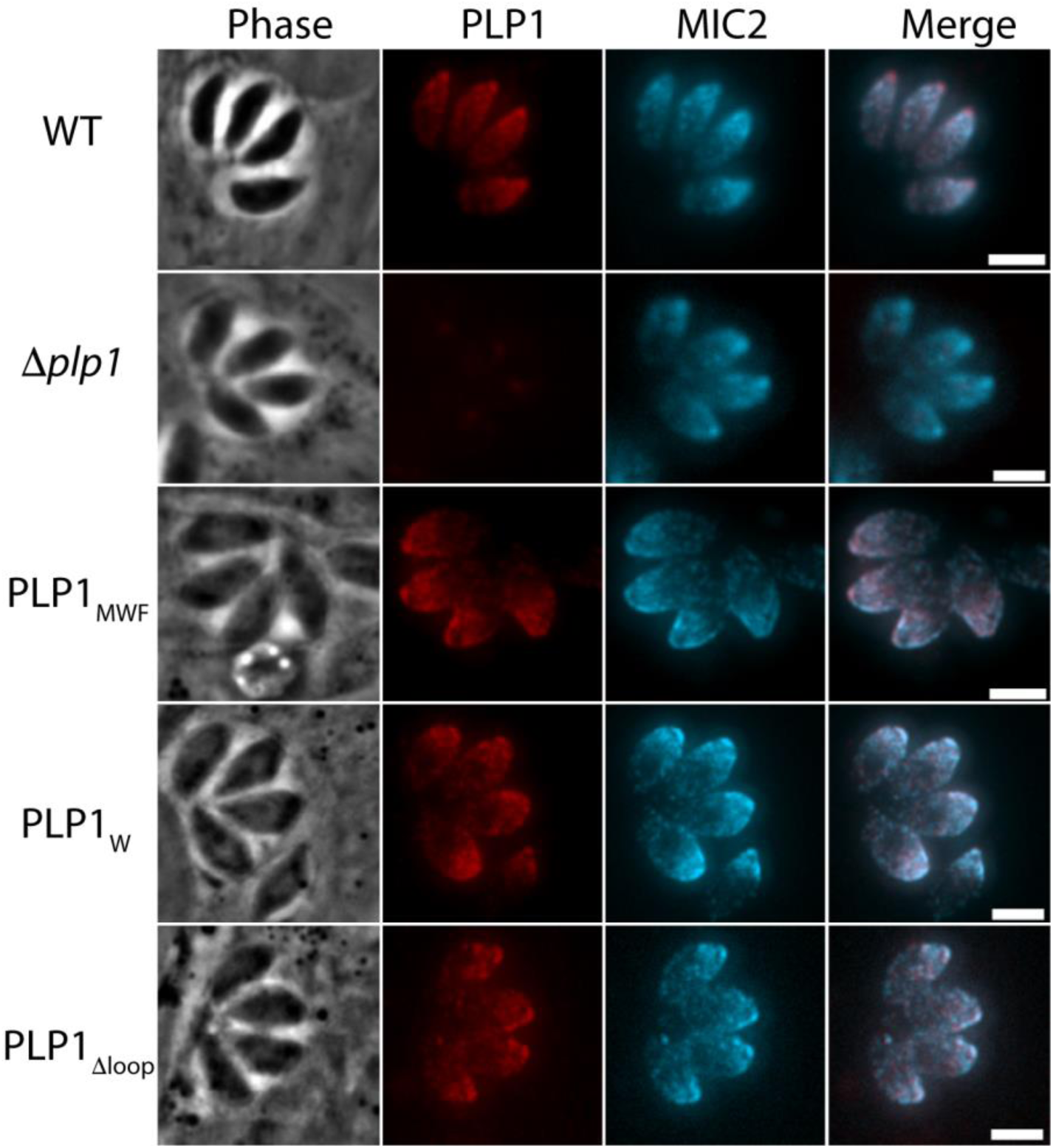
Indirect immunofluorescence of intracellular parasites with mouse anti-*Tg*MIC2 (cyan) and rabbit anti-*Tg*PLP1 (red) antibodies.

**Figure S5.**
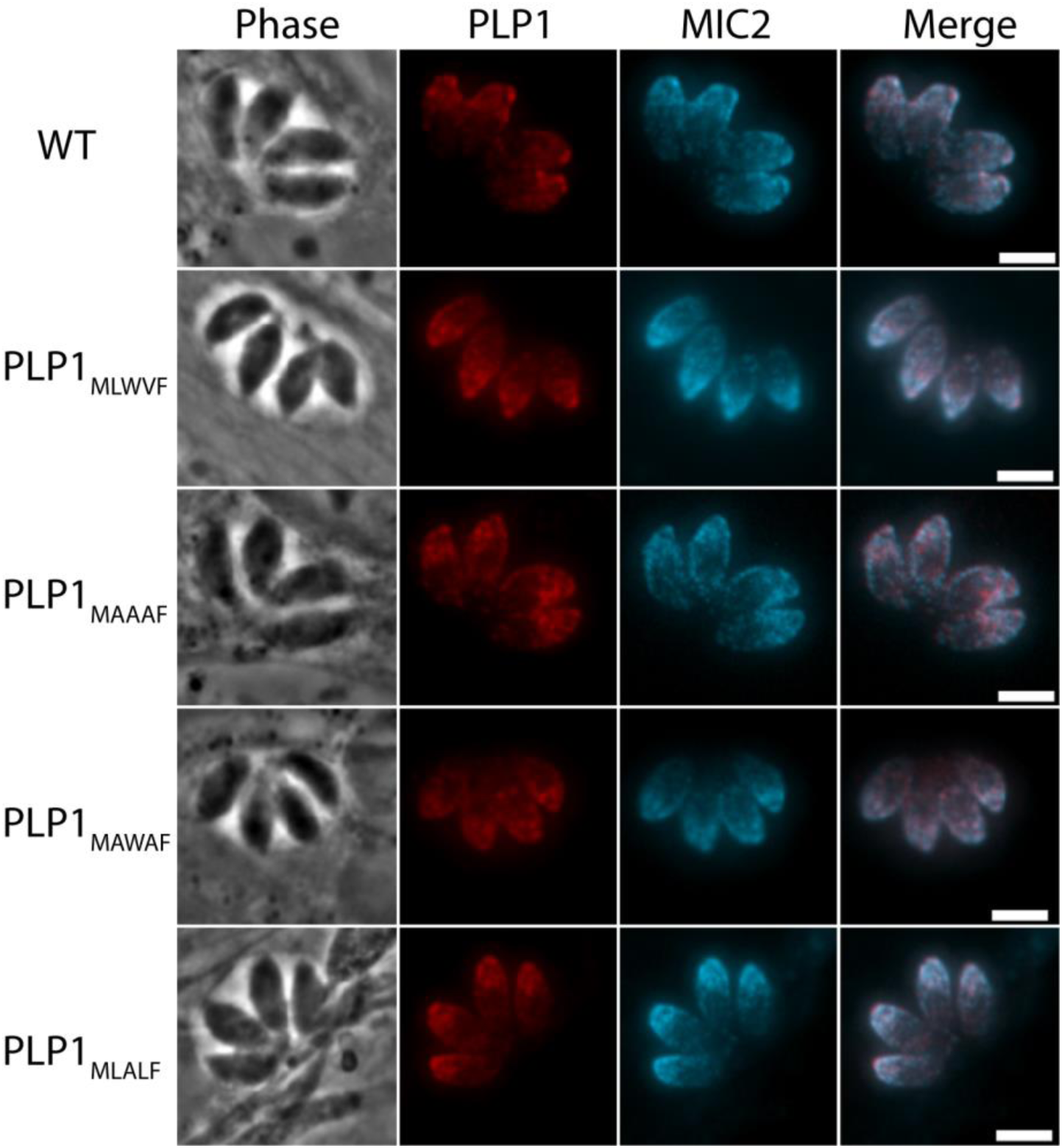
Indirect immunofluorescence of intracellular parasites with mouse anti-*Tg*MIC2 (cyan) and rabbit anti-*Tg*PLP1 (red) antibodies.

**Table S1.** Dali server search results for PDB entry 6D7A

| Number | PDB-chain | Dali-Z | % ID | Molecule Description |
| --- | --- | --- | --- | --- |
| 1 | 5ouo-A | 48.5 | 100 | Perforin-like protein 1 |
| 2 | 5vlp-A | 6.0 | 17 | Proprotein convertase subtilisin/kexin type 9 |
| 3 | 5a3k-A | 3.9 | 10 | Putative pteridine-dependent dioxygenase |

